# Individual, but not population asymmetries, are modulated by social environment and genotype in *Drosophila melanogaster*

**DOI:** 10.1101/694901

**Authors:** Elisabetta Versace, Matteo Caffini, Zach Werkhoven, Benjamin L. de Bivort

**Affiliations:** Department of Organismic and Evolutionary Biology and Center for Brain Science, Harvard University (USA); Department of Biological and Experimental Psychology, Queen Mary University of London (UK)

**Keywords:** Social behavior, individual behavior, behavioral asymmetries, individual-level lateralization, population-level lateralization, *Drosophila melanogaster*, DGRP, position, orientation, wing use, circling asymmetry

## Abstract

Theory predicts that social interactions can induce an alignment of behavioral asymmetries between individuals (i.e., population-level lateralization), but evidence for this effect is mixed. To understand how interaction with other individuals affects behavioral asymmetries, we systematically manipulated the social environment of *Drosophila melanogaster*, testing individual flies and dyads (female-male, female-female and male-male pairs). In these social contexts we measured individual and population asymmetries in individual behaviors (circling asymmetry, wing use) and dyadic behaviors (relative position and orientation between two flies) in five different genotypes. We reasoned that if coordination between individuals drives alignment of behavioral asymmetries, greater alignment at the population-level should be observed in social contexts compared to solitary individuals. We observed that the presence of other individuals influenced the behavior and position of flies but had unexpected effects with respect to individual and population asymmetries: individual-level asymmetries were strong and modulated by the social context but population-level asymmetries were mild or absent. Moreover, the strength of individual-level asymmetries differed between strains, but this was not the case for population-level asymmetries. These findings suggest that the degree of social interaction found in Drosophila is insufficient to drive population-level behavioral asymmetries.

## Introduction

Consistent left-right asymmetries in brain and behavior are widespread among animal species ^1–4^. For instance, the vast majority of people is right handed (a behavior controlled by the left hemisphere), and an advantage of using the left eye (right hemisphere) has been observed in agonistic interactions in chicks ^5^, lizards ^6^, toads^7^ and baboons ^8^, while in bees the use of left antennae enhances aggressive behavior ^9^ and the use of the right antenna is involved in social behavior ^10^. In these cases, when asymmetries are aligned on the same side in the majority of the population, we consider this population-level lateralization or directional asymmetry. In other cases, individuals exhibit strong and consistent preferences for one side, but these are not aligned at the population level and we define these cases individual-level lateralization or antisymmetry.

How the presence of other individuals influences these asymmetries at the individual and group level remains largely unknown. Mathematical models suggest that selective pressures associated with living in social contexts enhance the alignment of behavioral asymmetries (population-level lateralization), due to the advantages of coordinating between individuals ^11,12^. This hypothesis predicts population-level asymmetries in social contexts more than in individual contexts, for the contexts that occurred repeatedly in the course of evolution (for a recent review see ^13^). The growing evidence on the effect of the social context on asymmetric behavior, though, is mixed ^9,14^. Contexts that are expected to elicit coordinated social behavior are parent-offspring, female-male and agonistic interactions or coordinated group movements. Several works directly support this idea; for instance, gregarious species such as sheep coordinate motor biases within populations and maintain the same side bias across generations ^15^, in many species social interactions between mother and offspring are lateralized at the population level ^16,17^, and population asymmetries have been observed in mating and fighting in several species ^18–20^. In social insects such as honeybees, bumblebees and social stingless bees, a strong population bias in the use of antennae for olfactory learning has been observed, contrary to solitary bees that do not exhibit a population bias ^21^. This evidence suggests that population lateralization might have evolved under the pressure to coordinate between individuals. While this model focuses on explaining the alignment of population-level asymmetries, it does not predict a modulation of individual-level asymmetries as a function of the social context.

To date, individual level lateralization has been explained as the result of advantages that are mostly independent from social interactions ^13,22^, such as avoiding neural reduplication in small nervous systems, simultaneous and parallel processing of information, increased/faster/stronger motor control and cognitive specialization ^22,23^. Data from several taxa are consistent with such advantages, for instance: strongly lateralized chimpanzees are more effective at fishing termites using a stick ^24^, lateralized parrots are better at solving novel problems (string pulling and pebble-seed discrimination) than less lateralized parrots ^25^, lateralized antlions have advantages in learning ^26^, and lateralized locust perform better crossing a gap ^27^. To our knowledge, no studies have investigated whether individual-level asymmetry is modulated by social context, nor is there theory predicting effects one way or the other.

Population level asymmetries have been identified not only in eusocial or gregarious species, but also in interactions of “solitary” species. Some examples include aggressive/mating contexts in solitary mason bees ^9^, blowfly and tephritid flies ^19,20^, locusts during predator surveillance ^28^. There are also examples of population asymmetries in nominally solitary behaviors in solitary species, including: sensory asymmetries in nematodes ^29^, *Drosophila* larvae ^30^ and adults ^31^. It is not clear, though, whether these asymmetries are connected to social interactions: larvae and adult flies aggregate and interact during foraging, and locusts exhibit collective migration in their gregarious phase. Moreover, for sexually reproducing animals, it is difficult to rule out the possibility that population-level asymmetries are related to social interactions, since all such animals perform the social behavior of mating. An experimental approach in which the social context is manipulated and the effects on lateralization are monitored might clarify the role of the social context on lateralization.

Here we adopted this approach to understand how social context affects population and individual level lateralization. We use fruit flies (*Drosophila melanogaster*) to assess how individual-level and population-level asymmetries are affected by social context (two males, two females, one male and one female, one male or one female). As social behaviors, that theory predicts will be modulated by social context, we examined relative position and orientation between individuals. As individual behaviors, that are not expected to be strongly modulated by the social context, we examined circling asymmetry (clockwise/anticlockwise circulation in an arena) and preferential wing use.

The social asymmetry hypothesis predicts: (a) stronger population-level asymmetries in social versus individual contexts, (b) stronger population-level asymmetries in social versus individual behaviors, (c) a modulation of population-level asymmetries with the particular social context, with larger population-level asymmetries in male-female (courtship) and male-male (potentially aggressive rivalry) interactions versus female-female interactions. Lack of these patterns would argue against a causative role of social context in population asymmetries in *Drosophila*.

### Dyadic behaviors

In our investigation of dyadic behaviors, we focused on the relative position between flies, a trait that is lateralized at the population level in different species ^1^. In many vertebrate species, systematic asymmetries in eye use are accompanied by asymmetries in body position, with a preferential use of the left eye for monitoring conspecific in birds, fish and primates ^32,33^. Moreover, several mammals preferentially keep offspring on the left side ^16^. It has been suggested that this reflects an advantage of the right hemisphere in social monitoring ^33,34^. It is not clear, though, whether flies exhibit side biases in their position and orientation. For instance, the male/female position during courtship systematically differs between species. In most species, such as *D. melanogaster*, males court females from behind, while in few species males court on the front or side/back, or even circle around the female ^35–38^.

The analysis of social interaction networks between same sex flies ^39^ has shown an effect of sex and genotype in the duration and propensity for interaction. Looking at short interactions between virgin flies of both sexes, a modulation of social context and sex has been observed, with males showing a higher probability of orienting themselves toward females than toward males, and females showing few biases in their orientation to other flies ^40^. In the same work, interspecific differences between *Drosophila* species were documented, indicating genetic variation for orientation asymmetries. The available evidence suggests that the relative position adopted during interactions is genetically modulated, but a systematic investigation of genetic variation for this trait is still lacking.

### Individual behaviors

As individual behaviors that do not require the presence of another fly, we studied circling asymmetry (clockwise/anticlock-wise circulation in a circular arena) and preferential wing use. Asymmetries at the individual but not the population level have been found in the clockwise/anticlockwise walking preference of individual flies ^41–43^ or male/female dyads ^44,45^ but little is known on the influence of social context.

When not flying, flies use their wings in self grooming (a behavior that is more frequent in social contexts ^46,473^), courtship and aggression. Male flies extend and vibrate wings during courtship in species-specific fashion. For instance, *D. melanogaster* mainly vibrates one wing, *D. suzukii* one or two wings, whereas *D. biarmipes* flutters both wings ^37^. *D. melanogaster* males make a greater use of wings when they are located behind the female, likely to perform courtship song ^48^. Wing threat is used by both males and females in aggressive contexts directed towards both sexes ^49^. Because flies use wings in social contexts, a modulation of population-level asymmetries in wing use might occur due to evolutionary pressures for coordination. Wing use asymmetry in flies has been studied for several decades ^44,45^ with different outcomes but population-level lateralization has not been observed. Buchanan and colleagues ^42^ found evidence of individual side biases in wing folding (right or left wing closed on top of the other wing). This trait was not correlated to asymmetries in clock-anticlockwise circulation. While some authors ^44,45^ observed individual preferences in wing folding, another ^50^ failed to observe consistent individual preferences. Hence, previous results are not conclusive.

## Methods

### Data, code and materials

Raw data and scripts are available in Zenodo: 10.5281/zenodo.3268870. The scripts are also archived with a read-me at http://lab.debivort.org/social-asymmetries.

### *Drosophila* stocks and husbandry

We used five inbred lines of the *Drosophila* Genetic Reference Panel ^51^ from the Bloomington *Drosophila* Stock Center. Each line in this collection was started as an isofemale line derived from a single field collection in Raleigh (North Carolina) and then subjected to 20 generations of full-sibling mating, making most loci homozygous within lines ^52^. We tested lines belonging to different mitochondrial clades (2012): line 69 (clade III), line 136 (clade I), line 338 (clade VI), line 535 (clade I), line 796 (clade I). Flies were reared on cornmeal/dextrose fly food (made in the Harvard University BioLabs Fly Food Facility) in a single incubator at 25°C at 30–40% relative humidity with a 12:12-h light:dark cycle. Flies were tested between day 3 and 9 post-eclosion. Before the test, flies were housed in standard fly vials with 30-40 conspecifics of both sexes. Table 1 displays the number of individuals tested in each condition for each line.

**Table 1.** Number of individuals tested for each line and sex.

|  | ♀ | ♂ | ♀-♀ | ♂-♂ | ♀-♂ |
| --- | --- | --- | --- | --- | --- |
| **RAL-69** | 99 | 100 | 152 | 156 | 216 |
| **RAL-136** | 100 | 100 | 150 | 196 | 188 |
| **RAL-338** | 100 | 100 | 180 | 138 | 200 |
| **RAL-535** | 100 | 100 | 160 | 154 | 160 |
| **RAL-796** | 100 | 100 | 160 | 160 | 150 |

### Apparatus

Behavior was measured in circular acrylic arenas arrayed for simultaneous imaging of 15-20 arenas. See description of imaging set-ups in ^53^. Arenas were illuminated from below by an array of white LEDs (Knema), imaged with digital cameras (Pointgrey Blackfly 1.2 MP Mono GigE PoE) and recorded at a rate of 10 frames per second with Pointgrey Fly-Capture software for 60 minutes (36,000 frames). Arenas (2.54 cm in diameter, 1.6 mm deep) were fabricated in black acrylic using a laser cutter. Each arena’s transparent acrylic lid was lubricated with Sigmacote (Sigma) to prevent flies from walking on the underside of the lids. To illuminate the arenas uniformly, we placed a diffuser (two sheets of 3.2 mm-thick transparent acrylic roughened on both sides by orbital sanding) between the LED array and the arena array.

### Phenotypic assay

Before the test, flies were lightly anaesthetized with CO_2_ and placed in the arena with a brush. The trial started after an acclimation of 12-15 minutes and lasted 60 minutes. Flies were tested individually or in dyads that contained two individuals of the same line (but previously kept in separate vials): two females, two males, or one male and one female. Assays were run between 10:00 am and 8 pm. Since preliminary analyses showed that the hours of the day had at most a small effect size on all dependent variables (in line with the previous results ^54^), we excluded this variable from further analyses. For each frame, the x and y positions of each fly centroid and the angle of the wings with respect to the body midline were measured with Flytracker ^55^. The released version of the software smooths the computed angles between flies using a spatial kernel of three values from subsequent frames (with weights 1/4, 1/2, 1/4, respectively). When the fly position is expressed in polar coordinates, this produces an error for flies positioned at 180°. To fix this error we modified the code excluding the angle smoothing function.

### Analyses

For each individual animal and frame we analyzed individual behaviors and dyadic behaviors.

As individual behaviors, we measured circling asymmetry *C* and asymmetrical wing use *W*. To obtain *C* for each fly, we measured positive and negative deviations in trajectory (degrees) across subsequent frames (see Figure 1), using the angle between the previous trajectory (vector between time t_1_ and time t_2_) and the current trajectory (vector between time t_2_ and time t_3_) and calculated the ratio between the sum of positive (anticlockwise) and negative (clockwise) deviations (*c*) and the overall deviations (sum of the absolute value of all deviations).

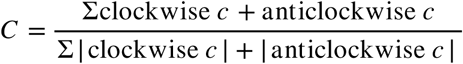

**Figure 1.**
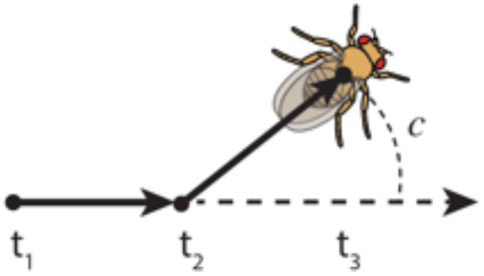
Vectors used to measure circling asymmetry, as the angle between the previous trajectory (vector between t_1_ and t_2_) and the current trajectory (vector between t_2_ and t_3_). Clockwise deviations are positive, anticlockwise deviations are negative.

Positive *C* indicates clockwise preferences, negative *C* indicates anticlockwise preferences. To quantify population-level and individual-level asymmetries we used *C* and |*C*| respectively, where significant departures from 0 indicate significant asymmetries.

To obtain *W* for each fly, we used a similar procedure and measured positive (right wing) and negative (left wing) degrees of wing opening in each frame and calculated the ratio between the sum of positive (left wing) and negative (right wing) opening degrees (*w*) and the overall wing opening (sum of the absolute value of all wing openings).

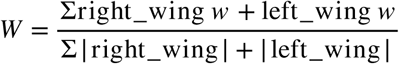

Positive W values indicate predominantly left wing use, negative values predominantly right wing use. To quantify population-level and individual-level asymmetries in wing use, we used *W* and |*W*| respectively, where significant departures from 0 indicate significant asymmetries.

### Dyadic behaviors

involved two subjects. For each frame, the spatial relationship between flies A and B was quantified by calculating: (a) the Distance between the centroids of the two flies [mm], (b) the relative Position as the angle in degrees between back-head vector of the focus fly and the segment vector the centroids of the two flies, (c) the Orientation, as the angle in degrees between the back-head vectors of the two flies. Of these three measures, the Position between the dyad partner fly (B) compared to the focal fly (A) can be lateralized. We scored the relative Position *P* of a fly (fly B in Figure 2b) as negative when it was located to the right of the focal fly, and positive when it was located to the left of the focal fly. Values close to +/-180° indicate that the focal fly is in front of the other, while values close to 0 indicate that the focal fly is located behind the other fly.

**Figure 2.**
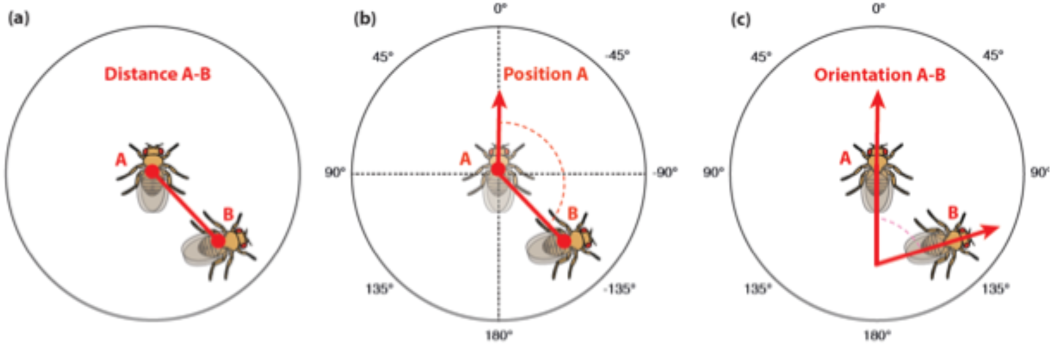
Social metrics calculated for dyads of flies. (a) Distance in mm between the centroids of the flies. (b) The Position of fly B is the angle between the vector trajectory of A and the vector between the centroids of A and B. When B is located to the right of A Position is negative, when B is located to the left of A Position is positive. (c) The relative Orientation between A and B is the angle between the facing vectors of the two flies. It ranges between 0°, when flies are parallel, and 180°, when flies are facing opposite directions.

### Statistics

We analyzed each variable using an ANOVA with social Context (two females, two males, one male and one female, single fly), Sex (male, female), Strain (RAL-69, RAL-136, RAL-338, RAL-535, RAL-796) and their interactions as independent variables. We set significant results with alpha level of 0.05 and considered to be biologically meaningful factors or interactions that not only were significant but also had a size effect medium or high as defined by ω^2^ (see ^56^) greater than 0.6. We used one-sample *t*-tests against the chance level (0) and Cohen’s d as effect size estimate to check for overall asymmetries. Statistical calculations were conducted with R 3.5.2.

## Results

### Individual behaviors: circling asymmetry and wing use

To investigate the effect of the social context and strain on the asymmetry of individual behaviors, we investigated clockwise/ anticlockwise circling asymmetry, and left/right asymmetry in wing positioning in individual males (M) and females (F) and in dyads with two females (FF), two males (MM), one female and one male (FM). If behavioral asymmetry is enhanced by social context, we would expect greater population-level asymmetries in dyads compared to individual flies, especially in FM and MM dyads, due to the evolutionary importance of mating and aggression. Moreover, we would not necessarily expect modulation of individual-level asymmetries.

### Population-level circling asymmetry

For each fly, strain and context we calculated the circling asymmetry index across the whole trial (60 minutes), which represents each animal’s tendency to turn clockwise or anticlockwise (this is measured as the population circling asymmetry index, *Cp*, which ranges from −1 to 1). All factors (Context, Sex and Strain) provided only a very small explanatory contribution to the overall variance and there was no significant difference between single flies and dyads (see Figure 3 and Supplementary Table 1). These results suggest that, in dyads, social context does not modulate population-level circling asymmetry.

**Figure 3.**
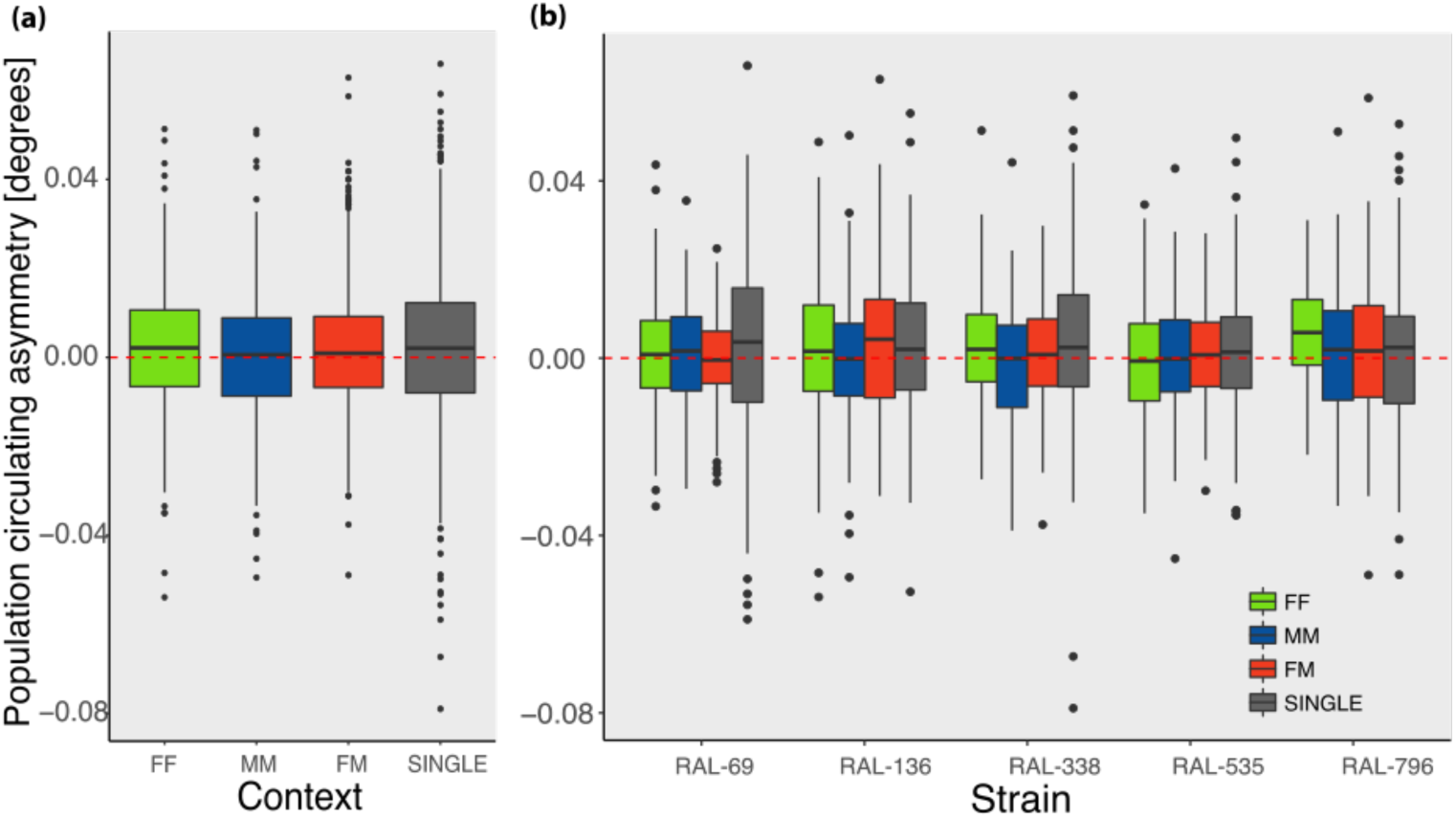
(a) Overall mean population circling asymmetry by Context (FF, MM, FM, SINGLE) and (b) by Context and Strain. The dashed line indicates the absence of population-level circling asymmetry. Here and elsewhere, boxes demarcate the interquartile range, thick horizontal line the median, whiskers minimum/maximum excluding the outliers, and point outside this range outliers, namely points greater/lower than 1.5 times the interquartile range).

We observed a very small but significant preference for the circling anticlockwise asymmetry (t_3518_=6.3, p=2e-10, Mean=0.0015, SD=0.014, d=0.11, 95% CI: 0.0010-0.0020; see Figure 3b). In line with this small effect, previous assays did not show population-level asymmetries ^41,42,44,57^. We did not observe any increase in population alignment in dyads compared to individual flies (see Supplementary Table 2), suggesting that the evolutionary pressure for alignment with conspecifics is absent or negligible in circling asymmetry in fruit flies.

### Individual-level circling asymmetry

For each fly, strain and context we calculated an absolute circling asymmetry index across the trial (60 minutes). This index ranges from 0 (no individual bias) to 1 (complete preference for either clockwise or anticlockwise circling asymmetry). Exploratory analyses showed a constant level of individual asymmetry during the test. We observed a significant main effect and high ω^2^ only for Context (F_1,3489_=123.521, p<2e-16, ω^2^=0.085, Figure 4a) and Context x Strain (F_12,3489_=35.358, p<2e-16, ω^2^=0.095, see Figure 4b), see Supplementary Table 3 for the complete results. Flies in the single context had lower individual asymmetry than flies in dyads (F_1,3499_=319.856, p<2e-16, ω^2^=0.075), and this factor (Dyad) was the main explanatory factor of the observed variance (Supplementary Table 4). The greater individual asymmetry in individual flies suggests that, similarly to what already documented in phototactic behavior in cockroaches ^58^, individual behavioral preferences in insects are modulated by whether a task is performed in isolation or in a group. Overall, looking at the individual circling asymmetry index vs. the absence of asymmetry (0) we observed strong individual-level lateralization (t_3518_=198, p<2e-16, Mean=0.340, SD=0.10, Cohen’s d=3.3, 95% CI=0.336, 0.343; see Figure 4a). This result confirms previous reports of strong individual preferences in locomotor behavior in flies tested individually ^41,42^.

**Figure 4.**
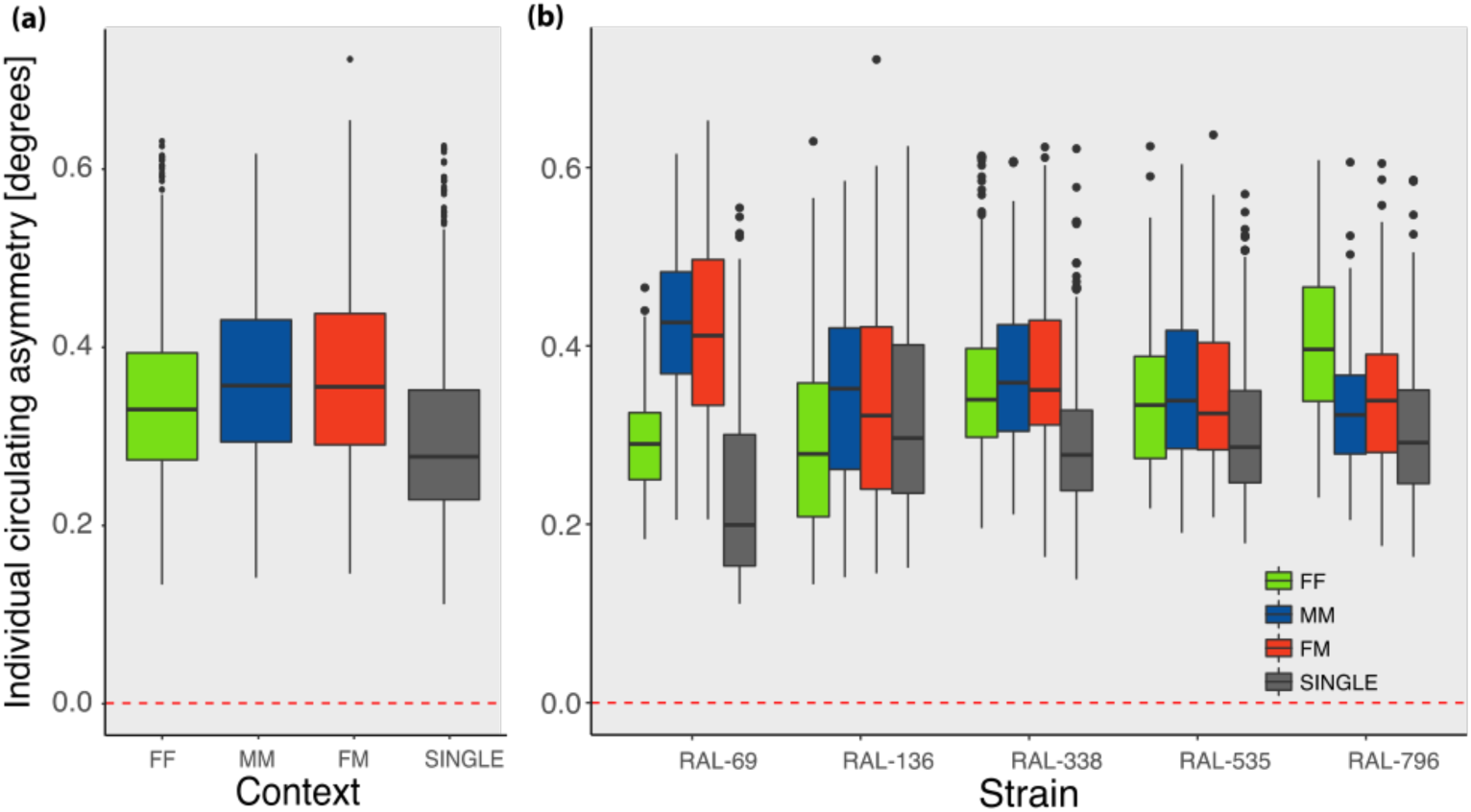
(a) Overall individual circling asymmetry index by Context (FF, MM, FM, SINGLE) and (b) by Context and Strain. The dashed line indicates the absence of individual-level circling asymmetry.

Contrary to our hypothesis, when comparing the data on population- and individual-level circling asymmetries, the effect of the social context (Context), Strain and their interaction appeared much stronger for individual than for population asymmetries. Moreover, while population-level circling asymmetries were small for all genotypes, individual asymmetries were strong for all genotypes (Figure 4b). This suggests that, in *Drosophila melanogaster*, circling asymmetry is modulated by the social context mostly at the individual level, whereas circling asymmetry is neither substantial or socially modulated at the population level. The presence of a strong individual level asymmetry and absence of population level asymmetry is clear comparing the histograms of the observed asymmetry indices (individual and population) and their respective expected distributions in the absence of asymmetries (Figure 5a,b). The social context increases individual but not population circling asymmetries in *Drosophila*. Interestingly, FM (female-male) dyads and MM (male-male) dyads, in which mating and aggression are expected at high frequencies, did not exhibit greater population asymmetries. Overall, our results on circling asymmetry appear inconsistent with a role of social context as a driver of population motor asymmetries, but indicate a novel role in modulating individual motor asymmetry.

### Population-level and individual-level wing use asymmetry

For each fly, strain and context we calculated the wing use asymmetry index for the whole trial (60 minutes). Exploratory analyses showed that this metric was stable across the experiment. For population-level lateralization there was no significant effect of Context, Sex, Strain or their interaction (Supplementary Table 5), nor a significant difference from the chance level (t_3518_ = −0.677, p = 0.498, Mean=-0.0007, SD=0.058). For individual level lateralization we observed significant effect with a very small effect size for Context, Sex, Strain, Condition x Strain, Condition x Sex x Strain (Supplementary Table 6).

### Position

The distributions of Positions (see Figure 2b) kept by individual flies of all tested genotypes in FF, MM and FM dyads (where 0° indicates a Position in the front of the focal fly) are shown in Figure 6. The Position of different strains and contexts is shown in Figure 7.

**Figure 6.**
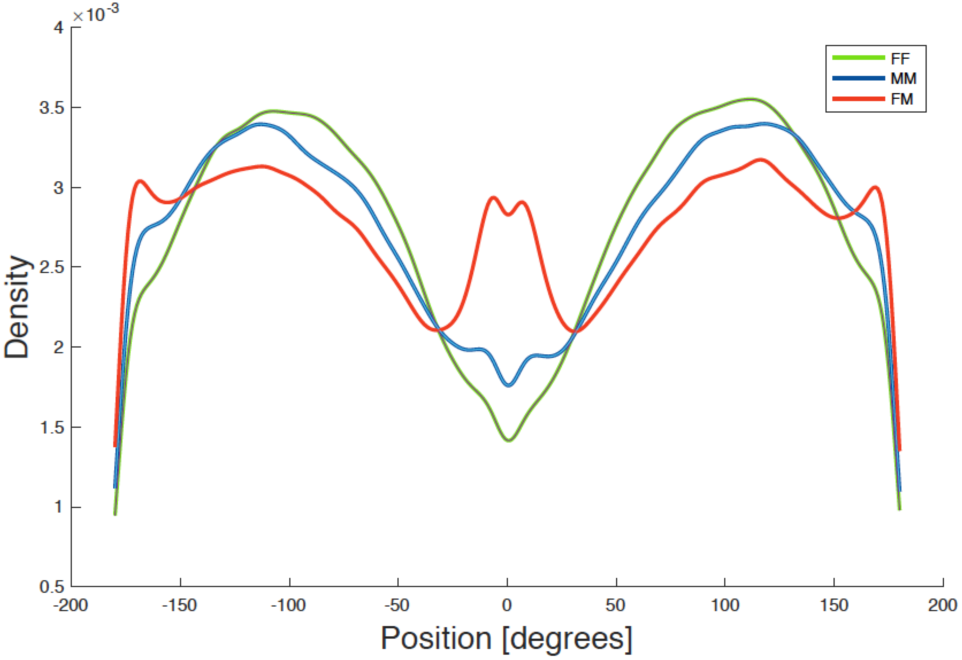
Kernel density distribution of the Position (in degrees) exhibited by the overall sample (all genotypes) during the test in each Context.

**Figure 7.**
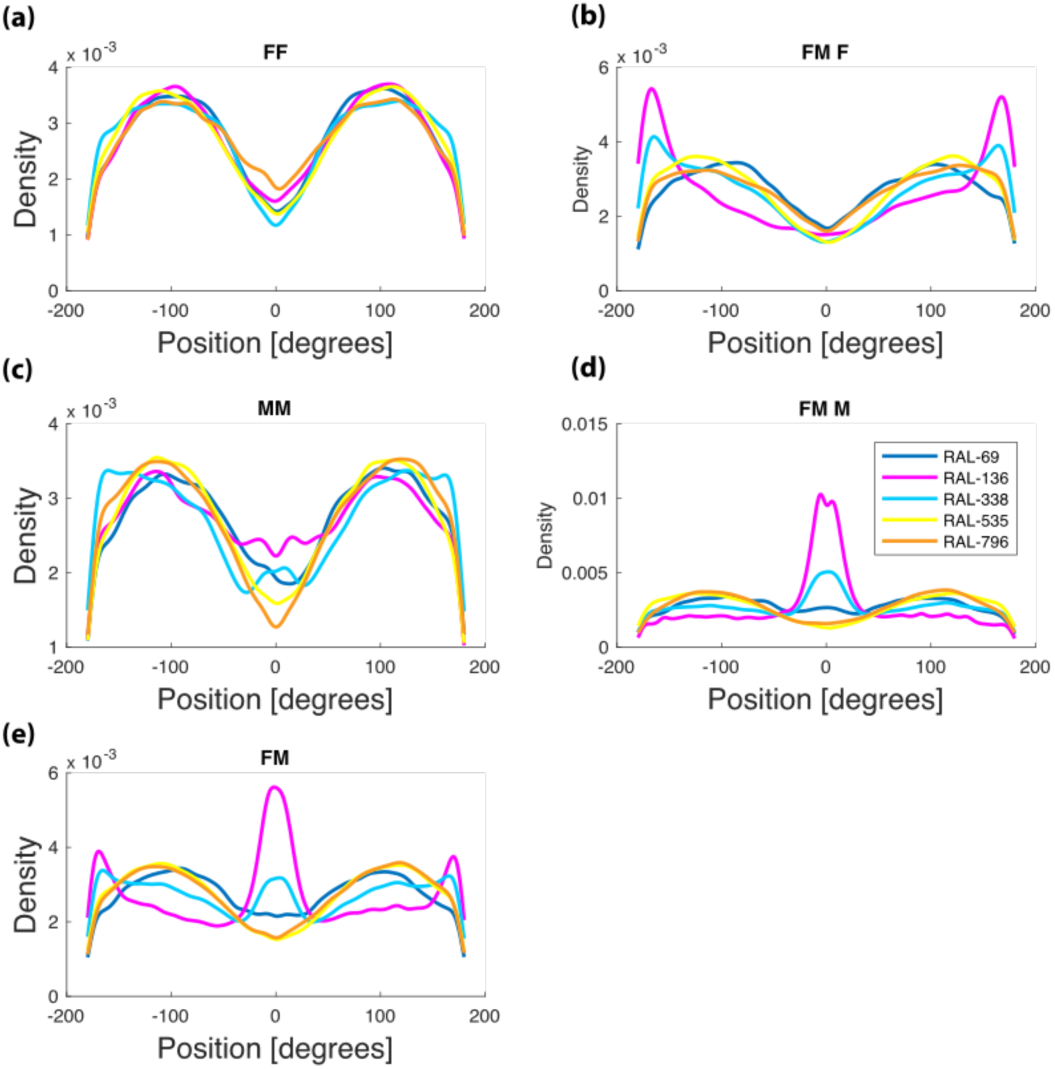
Kernel density distribution of Position (in degrees) kept during the test in each Strain, Context and Sex: (a) female-female dyads by strain, (b) females of female-male dyads by strain, (c) male-male dyads by strain, (d) males of female-male dyads by strain, (e) female-male dyads.

Overall, and in each strain separately, same sex dyads (FF and MM) preferentially chose oblique Positions to the other fly (Position ∼ 100°). In FM dyads there were additional modes at Position 0° and 180°, which correspond to males positioning females ahead of them and females positioning males behind them. In the genotypes RAL-136 and RAL-338, these relationships were particularly strong, but, surprisingly, RAL-535 and RAL-796 did not show these courtship-associated modes in Position (Figure 7). These observations suggest a strong interaction between genotype and social context. No side bias in the Position of flies was present in any social context or strain. Hence, in spite of a clear front-back modulation of the Position, the right-left dimension was not modulated by social context or strain.

The joint distributions of Position and Distance between focal flies of all tested genotypes and their dyad partners are shown for each context in the polar heatmaps of Figure 8a. Flies in dyads of the same sex keep their partner to the side and behind them in two ring-shaped regions of density, one on each side (note that this particular pattern likely arises through an interaction of the flies’ behavior and the arena geometry). The main interactions at close Distance are seen in female-male dyads and, to a less extent, male-male dyads. Large differences are observed between genotypes, especially in male-male and female-male dyads (Figure 8b). These plots confirm the absence of asymmetrical positioning in flies in all social contexts and at the same time the existence of standing genetic variation for other aspects related to the position of flies in different social contexts.

**Figure 8.**
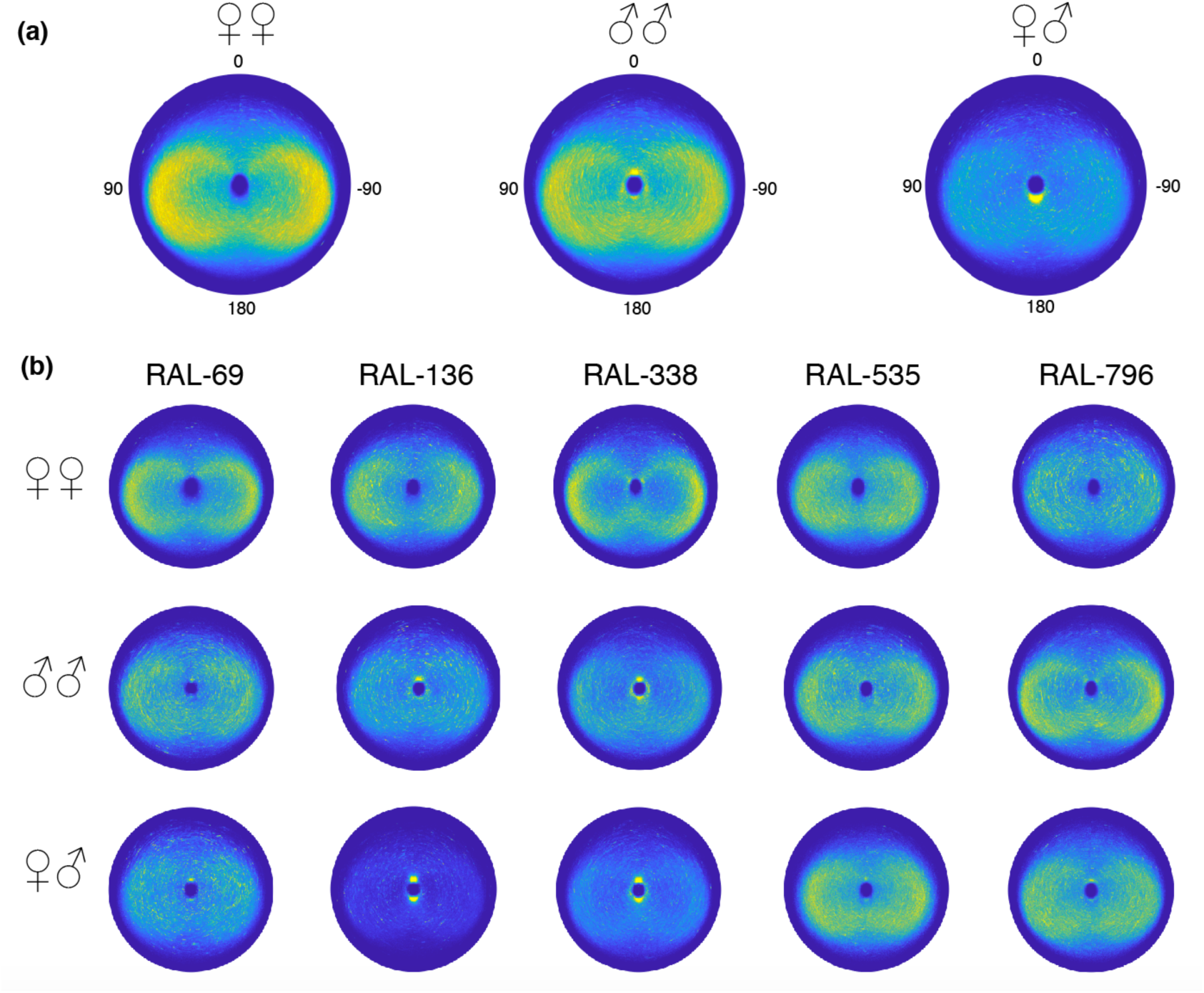
Heatmap in polar coordinates, showing the joint distribution of Position and Distance from the focal fly in each Context: (a) female-female, male-male and female-male dyads overall (b) female-female, male-male and female-male dyads for each strain.

Although not directly relevant for investigating the role of the social context on behavioral asymmetries, we analyzed the effect of Context and Strain on scalar behavioral measures that cannot be lateralized: Velocity, Distance and Orientation. The analysis of velocity, a measure of general activity, is presented in Supplementary materials 1. Briefly, we observed a strong effect of social Context (slowest speed in FM dyads, faster in single flies) and Strain (Figure S1.1 and S1.2), and a tendency to increase in velocity over the course of the experiment (Figure S1.3.)

The Distance between larvae and adult individuals is used as a measure of social interaction in *Drosophila* ^62–68^ as well as in other species ^69–73^. An effect of strain and sex has been previously found on Distance in the DGRP lines ^62^. In our experiments, Distance between flies was clearly modulated by the social environment, genotype and their interaction (see Supplementary Materials 2: Distance). Briefly, we observed a strong effect of Context, Strain and their interaction (Figures S2.1 and S2.2). Flies in the FM and MM contexts stayed closer than expected by chance and RAL-136 and RAL-338 had a peak around 3 mm, a distance characteristic of courtship.

## Discussion

To date, evidence that social interactions induce and sustain an alignment of behavioral asymmetries between individuals (population-level lateralization) is inconclusive. To shed light to this issue we analyzed individual and population-level asymmetries in *Drosophila melanogaster* in different social contexts. We systematically manipulated the social environment, observing the spontaneous behavior of individual flies and dyads (male-female, female-female, male-male). We looked at asymmetries in individual behaviors (circling asymmetry and wing use, that do not require partners) and dyadic behaviors (relative position of the partner fly) in five different isogenic strains ^62^. The hypothesis that the social context drives the alignment in behavioral asymmetries ^11,12^ predicts a potential increase of population-level lateralization with higher social engagement, likely in an FM > MM > FF trend.

Contrary to this prediction, we observed very little lateralization at the population level in any social context. Instead, individual asymmetries, circling asymmetry in particular, went up in dyads compared to single-fly experiments. Moreover, individual lateralization was generally highest in female-male and male-male dyads (though this was genotype-dependent). These results suggest that the social context affects lateralization in flies, but only at the individual level. The presence of individual-level asymmetries suggests that the absence of population alignments is not due to detrimental effects of individual asymmetries. On the contrary, the large differences between genotypes suggest that variability in individual lateralization is a trait not subject negative selection. Other aspects related to the position of flies and their interaction, such as distance and velocity, were strongly modulated by the genotype and social context, showing that the FM, FF, MM and individual test conditions have biologically significant differences.

Previous reports showed neuroanatomical and sensorimotor population-level asymmetries in this species. For instance, *D. melanogaster* exhibits a strong population asymmetry in the location of the asymmetrical body in the right part of the brain ^74,75^. This structure is associated with enhanced memory. Moreover, sensory signals coming from the left antenna contribute more to odour tracking than those from the right in adult flies ^31^ and in larvae the right olfactory system performs significantly better chemotaxis than the left ^30^. Besides these asymmetrical traits, flies exhibit visceral asymmetries that include the S shape of the stomach, the left-handed looping of the testes around the vas deferens and the clockwise rotation of the genital plate (see ^74^ for a review). To which extent specific population-level asymmetries derive and are maintained by the advantages of coordinating between individuals is an empirical question. For asymmetries of the viscera that emerge during embryonic development (primary asymmetries) and whose disruption produces pathological conditions ^76^, it seems unlikely that the need for social coordination was the primary evolutionary force in place.

Although *D. melanogaster* is considered a solitary species (because flies do not form cohesive social groups nor cooperate in rearing offspring ^77^), social habits are nevertheless central ^39^. For instance, *Drosophila* larvae cooperate in burrowing to dig more effectively ^78,79^ and attract each other through pheromones ^80^. Moreover, adult flies aggregate on food and oviposition sites, select food patches based on the presence of conspecifics ^81,82^, use collective behavioral responses to avoid aversive cues ^83^, interact when competing for resources ^49^ or courtship and mating (reviewed in ^36^), and exhibit social learning ^84,85^. In spite of this, in our assays we did not observe any behavioral alignment at the population level, even in traits such as the right-left position in a dyad, which are lateralized at the population level in other species ^5,15–17,59,60^.

The absence of population-level lateralization we documented is compatible with different scenarios. One possibility is that population-level lateralization of sensorimotor behavior previously documented in *Drosophila* is driven by social pressures while the behaviors we have investigated are not, and that for this reason we have not observed population asymmetries. It seems unlikely, though, that more advantages are available for sensorimotor population alignment in chemosensory tracking than in male-female interactions. So, we think that this scenario is unlikely and incompatible with the lack of response to selection of population asymmetries. Further studies should clarify the developmental origin of these asymmetries, and whether they are related to the primary visceral asymmetries that emerge in early stages of embryonic development.

A possibility is that a selective pressure actively opposes the motor alignment between individuals in *Drosophila* either because the alignment would directly decrease flies’ fitness or because of pleiotropic effects. This possibility is indirectly supported by several lines of evidence. First, among the many traits that have been subject to artificial selection in *Drosophila*, population-level asymmetries (also called directional asymmetries) have been suggested to be the only traits that do not respond to selection ^86,87^. This idea is based on selection studies on morphological asymmetries in *Drosophila subobscura* ^88^, directional wing-folding ^89^ and asymmetry for eye size ^86^ in *D. melanogaster*, wing-folding and Y-maze choice preference in *D. melanogaster* and *D. paulistorum* ^44^, as well as by lack of mutational variance for population-level alignments ^90^.

In contrast with population-level asymmetry, genetic variability for individual-level asymmetry has been consistently observed. Individual-level asymmetries in different *Drosophila* species have been observed for circling behavior, tapping and wing extension ^44^. More recently, Ayroles and colleagues ^41^ have documented individual biases in Y-maze choices and circling behavior in *D. melanogaster*. They observed that the average magnitude of the left-right locomotor bias is heritable, contrary to the average magnitude of the bias. Although within-genotype variability of individually lateralized behavior in *Drosophila* is increased by environmental variability, the effect of genotype and genotype x environment interaction has a greater impact on the extent of individual side preferences ^91^. While it is not easy to select for consistent directionality, random asymmetry (the magnitude of difference between right and left bias) responds readily to selection in *Drosophila* ^92,93^.

Using a large sample (above 3500 individuals) and an automated precisely quantitative approach ^55^, we have clarified that individual-level but not population-level behavioral asymmetries are modulated by the social context in a genotype-dependent way in *Drosophila*. In the light of the available evidence, our findings suggest that there is no genetic variability for population-level behavioral lateralization in *Drosophila*, although individual asymmetries are not selected against. Moreover, we saw no evidence that the strength of social interactions drove population-level lateralization in either individual or dyadic locomotor behaviors. This argues against the generality of the social lateralization hypothesis.

## Contributions

EV conceived the experiments, ran the experiments, analyzed the data, wrote the manuscript; MC provided software tools, helped with data analyses and revised the manuscript; ZW ran the experiments, analyzed data, and revised the manuscript; BdB conceived the experiments, coordinated the study, analyzed data, and revised the manuscript. All authors gave final approval for publication and are accountable for all aspects of their work.

## Acknowledgements

We thank the Research Computing of FAS and the Neuroimaging Core (Ed Soucy, Joel Greenwood and Adam Bercu) at Harvard University for their excellent technical support. EV was funded by the Harvard Mind Brain and Behavior Faculty Award. BdB was supported by a Sloan Research Fellowship, a Klingenstein-Simons Fellowship Award, a Smith Family Odyssey Award, and the National Science Foundation under grant no. IOS-1557913.

## Supplementary materials

**Supplementary Table 1.** Results of the ANOVA and ω^2^ values on population-level circulating asymmetry for all conditions with Context, Sex and Strain as independent variables. Bold cells indicate significant results.

| | Df | F | p | $\omega^2$ |
| --- | --- | --- | --- | --- |
| **Context** | **3** | **3.624** | **0.0125** | 0.00222 |
| Sex | 1 | 0.423 | 0.5156 | -0.0002 |
| Strain | 4 | 0.783 | 0.5363 | -0.0002 |
| Context x Sex | 1 | 0.250 | 0.6170 | -0.0002 |
| **Context x Strain** | **12** | **2.093** | **0.0146** | 0.0037 |
| **Sex x Strain** | **4** | **3.271** | **0.0110** | 0.0025 |
| Context x Sex x Strain | 4 | 0.217 | 0.9291 | -0.0008 |
| Residuals | 3489 |  |  |  |

**Supplementary Table 2.** Result of the ANOVA and ω^2^ values on individual-level circulating asymmetry for all conditions with Dyad (Single, Dyad), Sex and Strain as independent variables. Bold cells indicate significant results.

| | Df | F | p | $\omega^2$ |
| --- | --- | --- | --- | --- |
| Dyad | 1 | 1.548 | 0.2135 | <0.001 |
| Sex | 1 | 2.301 | 0.1294 | 0.0004 |
| Strain | 4 | 0.724 | 0.5757 | -0.003 |
| DyadxSex | 1 | 3.751 | 0.0529 | 0.0008 |
| **Dyad x Strain** | **4** | **2.656** | **0.0313** | 0.0019 |
| **Sex x Strain** | **4** | **2.412** | **0.0470** | 0.0016 |
| Dyad x Sex x Strain | 4 | 0.372 | 0.8288 | -0.0007 |
| Residuals | 3499 |  |  |  |

**Supplementary Table 3.**
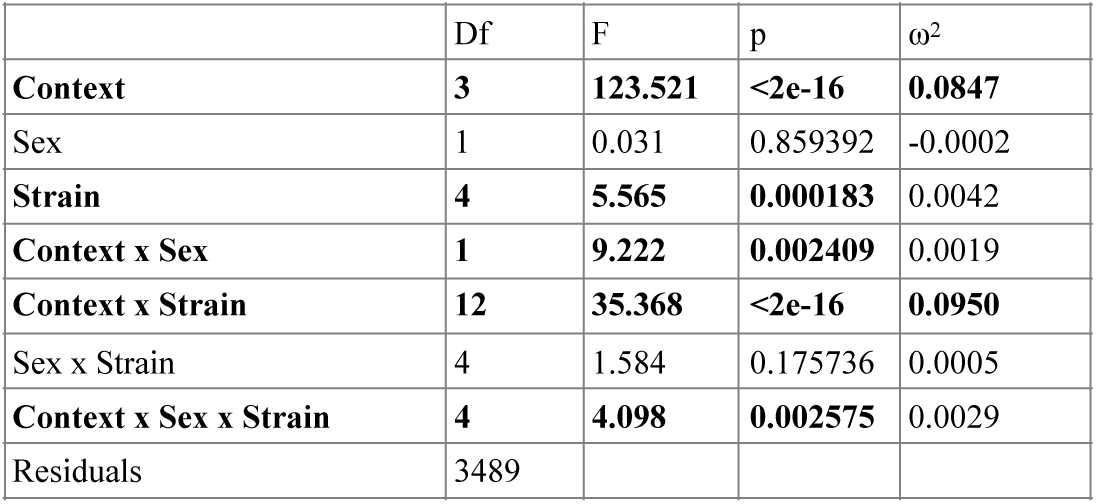
Results of the ANOVA and ω^2^ values on individual-level circulating asymmetry for all conditions with Context, Sex and Strain as independent variables. Bold cells indicate significant results.

**Supplementary Table 4.** Results of the ANOVA and ω^2^ values on individual-level circulating asymmetry for all conditions with Dyad, Sex and Strain as independent variables. Bold cells indicate significant results.

| | Df | F | p | $\omega^2$ |
| --- | --- | --- | --- | --- |
| **Dyad** | **1** | **319.856** | **<2e-16** | **0.0753** |
| **Sex** | **1** | **12.705** | **0.000370** | 0.0028 |
| **Strain** | **4** | **5.184** | **0.000366** | 0.0040 |
| **Dyad x Sex** | **1** | **24.439** | **8.03e-07** | 0.0055 |
| **Dyad x Strain** | **4** | **43.405** | **<2e-16** | 0.0401 |
| **Sex x Strain** | **4** | **30.146** | **<2e-16** | 0.0275 |
| **Dyad x Sex x Strain** | **4** | **15.311** | **2.04e-12** | 0.0135 |
| Residuals | 3499 |  |  |  |

**Supplementary Table 5.** Results of the ANOVA on wing use population-level asymmetry for all conditions with Dyad, Sex and Strain as independent variables.

| | Df | F | p | $\omega^2$ |
| --- | --- | --- | --- | --- |
| Context | 3 | 0.971 | 0.406 | 0.000 |
| Sex | 1 | 0.161 | 0.688 | 0.000 |
| Strain | 4 | 0.255 | 0.907 | -0.001 |
| Context x Sex | 1 | 0.004 | 0.948 | 0.000 |
| Context x Strain | 12 | 0.537 | 0.892 | -0.002 |
| Sex x Strain | 4 | 0.774 | 0.542 | 0.000 |
| Context x Sex x Strain | 4 | 0.664 | 0.617 | 0.000 |
| Residuals | 3489 |  |  |  |

**Supplementary Table 6.**
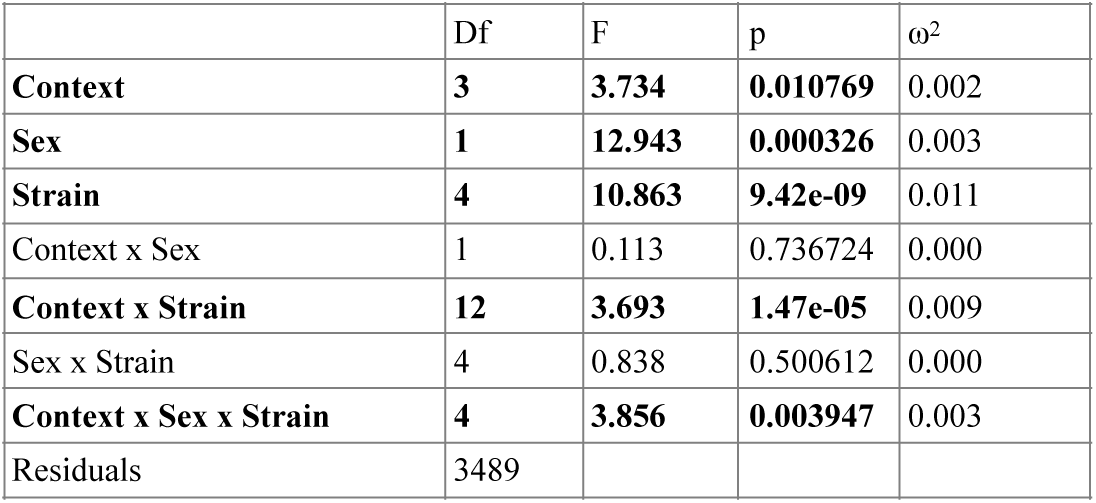
Results of the ANOVA on wing use individual-level asymmetry for all conditions with Dyad, Sex and Strain as independent variables. Bold cells indicate significant results.

| | Df | F | p | $\omega^2$ |
| --- | --- | --- | --- | --- |
| **Context** | **3** | **3.734** | **0.010769** | 0.002 |
| **Sex** | **1** | **12.943** | **0.000326** | 0.003 |
| **Strain** | **4** | **10.863** | **9.42e-09** | 0.011 |
| Context x Sex | 1 | 0.113 | 0.736724 | 0.000 |
| **Context x Strain** | **12** | **3.693** | **1.47e-05** | 0.009 |
| Sex x Strain | 4 | 0.838 | 0.500612 | 0.000 |
| **Context x Sex x Strain** | **4** | **3.856** | **0.003947** | 0.003 |
| Residuals | 3489 |  |  |  |

## Supplementary Results

### 1 Velocity

As a proxy for general activity in the testing arena, we analyzed the average velocity of flies. When comparing all contexts (FF, MM, FM and single flies) we observed a significant effect and a high ω^2^ for Context (F_3,3489_=168.932, p<2e-16, ω^2^=0.10 see Figure S1.1a) and Strain (F_4,3489_=176, p<2e-16, ω^2^=0.14, see Figure S1.1b), while the other significant main effects and interactions had a small explanatory power, see Fig. 1c and Table S1.1 for the complete results.

Because Context includes dyads and single tested flies, we further explored the difference between these two nested variables. A post-hoc ANOVA showed that flies in the single context moved faster than dyads (see Figure S1.2a): F_1,3499_=487.41, p<2e-16, ω^2^=0.09, see Table S1.2 for the complete results. In contrast, an ANOVA on the dyadic contexts showed that the significant difference between contexts explained a small fraction of the variance (F_2,2500_=8.621, p<0.001, ω^2^=0.005), and that Strain was by far the strongest explanatory factor (F_4,2500_=118.946, p<2e-16, ω^2^=0.149), see Table S1.3 for the complete results. This is consistent with the finding that genotype had a bigger effect on the mean and variance of several behavioral measures than environmental treatment or the interaction of treatment and genotype ^1^. Locomotion faster in individual flies than in dyads with different sex had been previously showed ^2^, while it was not previously reported that same sex dyads moved slower than individual flies. This shows that the slower speed observed in dyads in not specifically due to intersexual behaviors.

**Figure S1.1.**
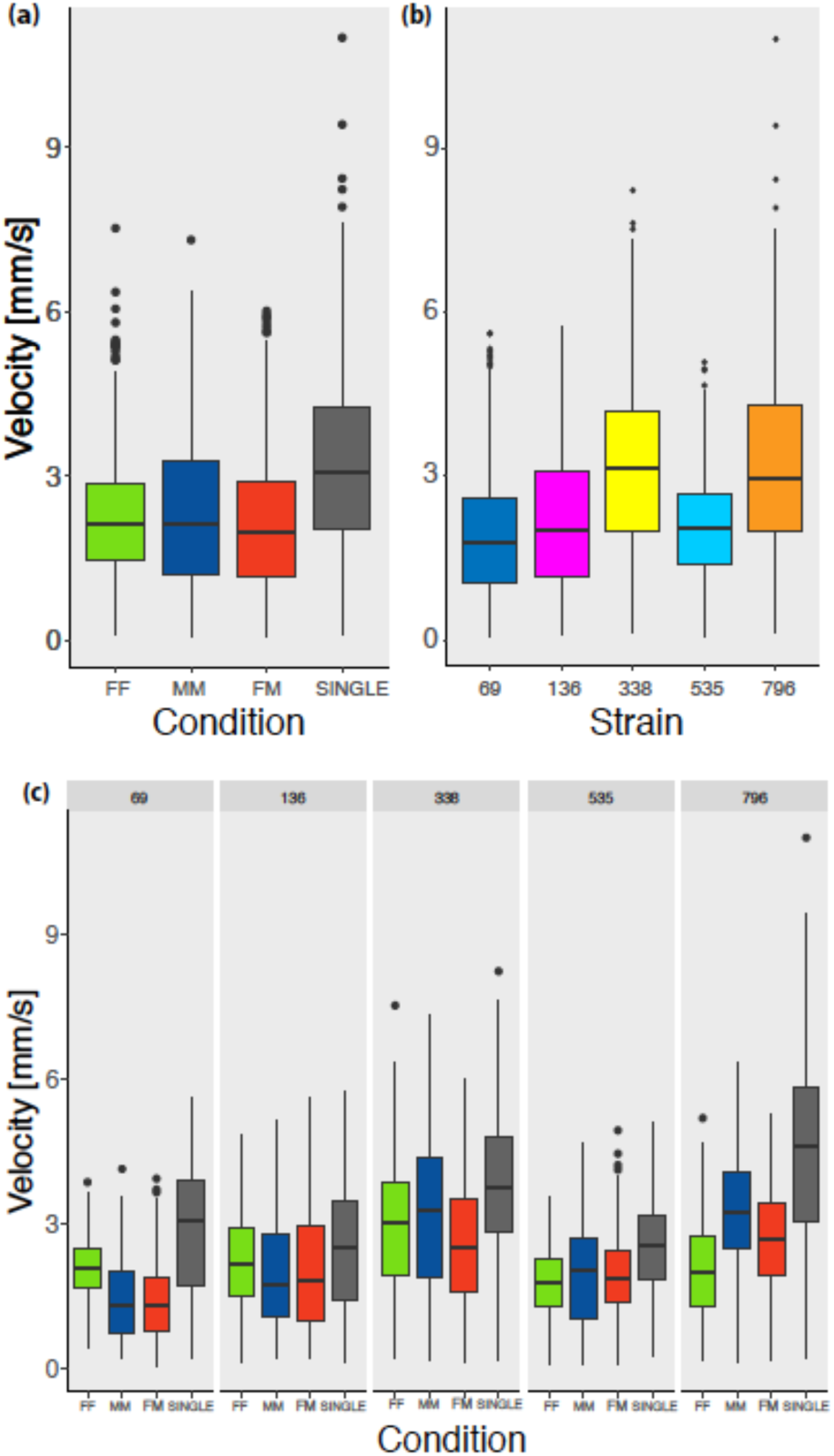
(a) Overall mean velocity by Context: dyads of two females (FF), two males (MM), one male and one female (FM), individual flies (SINGLE); (b) Overall mean velocity by Strain (c) Overall mean velocity by Context and Strain.

**Table S1.1.** Results of ANOVA and ω^2^ values on the variable Velocity for all conditions with Context, Sex and Strain as independent variables. Bold cells indicate significant results or a large portion of the variance explained by that factor.

| | Df | F | p | $\omega^2$ |
| --- | --- | --- | --- | --- |
| Context | 3 | 168.932 | <2e-16 | 0.1004 |
| Sex | 1 | 21.267 | 4.14e-06 | 0.0040 |
| Strain | 4 | 176.111 | <2e-16 | 0.1397 |
| Context x Sex | 1 | 14.910 | 0.000115 | 0.0028 |
| Context x Strain | 12 | 21.461 | <2e-16 | 0.0489 |
| Sex x Strain | 4 | 2.162 | 0.070729 | 0.0009 |
| Context x Sex x Strain | 4 | 2.856 | 0.022328 | 0.0015 |
| Residuals | 3489 |  |  |  |

**Table S1.2.** Results of the ANOVA and ω^2^ values on the variable Velocity for all conditions with Dyad, Sex and Strain as independent variables. Bold cells indicate significant results or a large portion of the variance explained by that factor.

| | Df | F | p | $\omega^2$ |
| --- | --- | --- | --- | --- |
| Dyad | 1 | 487.409 | <2e-16 | 0.0979 |
| Sex | 1 | 21.317 | 4.03e-06 | 0.0041 |
| Strain | 4 | 175.542 | <2e-16 | 0.1406 |
| Dyad x Sex | 1 | 14.420 | 0.000149 | 0.0027 |
| Dyad x Strain | 4 | 30.585 | <2e-16 | 0.0238 |
| Sex x Strain | 4 | 20.658 | <2e-16 | 0.0158 |
| Dyad x Sex x Strain | 4 | 9.165 | 2.30e-07 | 0.0066 |
| Residuals | 3499 |  |  |  |

**Table S1.3.** Results of the ANOVA and ω^2^ values on the variable Velocity for dyadic contexts with Context, Sex and Strain as independent variables. Bold cells indicate significant results or a large portion of the variance explained by that factor.

| | Df | F | p | $\omega^2$ |
| --- | --- | --- | --- | --- |
| Context | 2 | 8.621 | 0.000186 | 0.0048 |
| Sex | 1 | 0.192 | 0.661174 | -0.0003 |
| Strain | 4 | 118.946 | <2e-16 | 0.1486 |
| Context x Strain | 8 | 19.677 | <2e-16 | 0.0470 |
| Sex x Strain | 4 | 5.761 | 0.000130 |  |
| Residuals | 2500 |  |  |  |

A post-hoc ANOVA on the single tested flies showed that the main source of variability in Velocity was Strain (F_4,2500_=119, p<2e-16, ω^2^=0.23), while Sex had a significant effect but explained little variance (F_1,989_=3.9e-07, ω^2^=0.019) and the interaction Sex x Strain was not significant. See Table S1.4 for the complete results.

**Table S1.4.** Results of the ANOVA and ω^2^ values on the variable Velocity for the single flies with Sex and Strain as independent variables. Bold cells indicate significant results or a large portion of the variance explained by that factor.

| | Df | F | p | $\omega^2$ |
| --- | --- | --- | --- | --- |
| **Sex** | **1** | **26.094** | **3.9e-07** | 0.0190 |
| **Strain** | **4** | **76.846** | **<2e-16** | **0.2292** |
| Sex x Strain | 4 | 0.085 | 0.987 | -0.00276 |
| Residuals | 989 |  |  |  |

Overall, single flies consistently moved faster than flies in dyads (Figures S1.1a, S1.1c) with strong modulation by genotype, while the sex composition and social context had limited effects. Genetic variation for locomotor activity has been documented in the DGRP collection ^1,3^, including some of the strains investigated here, although other studies failed to observe significant differences between strains when comparing same sex groups ^4^. Here, we observe a significant difference between strains with a large portion of the variance explained by the factor Strain: F_3,380_=51.3, p<2e-16, ω^2^=0.282, see Figure S1.2).

Considering the number of turns in a Y-maze reported by Ayroles and colleagues ^3^ as a proxy for velocity (assuming that flies circulated in the maze using the entire space), we observe a significant difference between strains with a large portion of the variance explained by the factor Strain, similarly to what we found here: F_3,380_=51.28, p<2e-16, omega-squared=0.282, see Supplementary Figure S1.2).

Looking at time courses of Velocity for each Context and Sex (Figure S1.3), we observed that strains RAL-69, RAL-136 and RAL-535 showed roughly stable patterns over the course of each experiment, while RAL-338 and RAL-796 exhibited an increase of velocity in time, with the exception of females RAL-796 in FM dyads. An increase of locomotor activity due to starvation had been previously well described ^5–7^. It is likely that the increase of activity observed in some genotypes was a response to food deprivation, given that no food was provided in the test arena. It is clear that velocity trajectories across the experiment were modulated by both genotype and social context.

**Figure S1.2.**
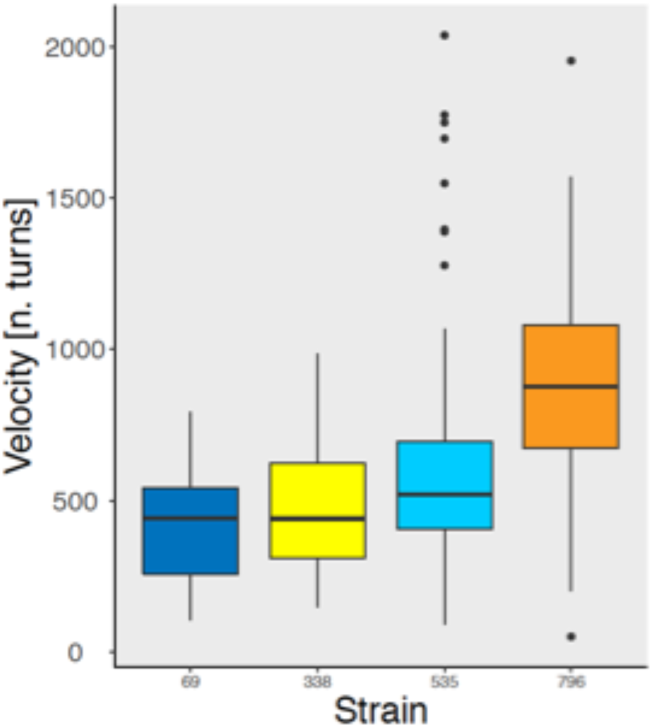
Velocity measured as number of turns in a Y-maze by Strain during two hours of observation (data from ^3^). The sample size is RAL-69=111, RAL-338=62, RAL-535=110, RAL-796=101.

**Figure S1.3.**
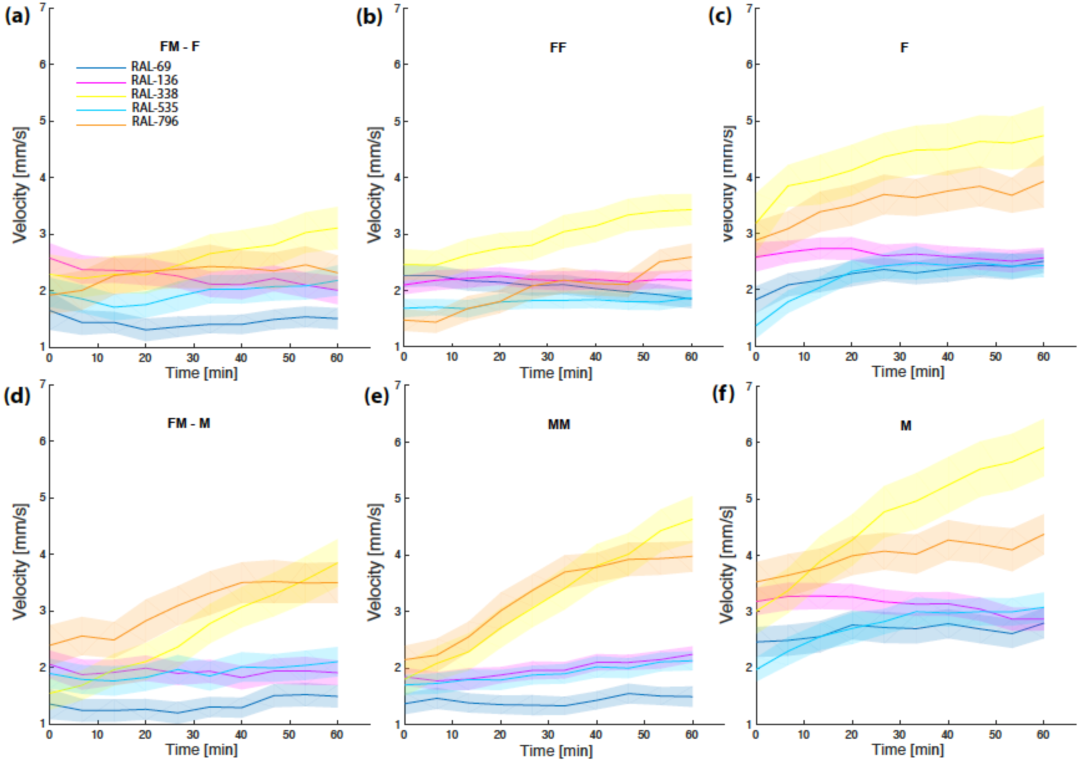
Velocity vs. time for different Strains, by Context and Sex. (a) Females in FM dyads, (b) females in FF dyads, (c) females tested individually, (d) females in FM dyads, (e) males in MM dyads, (f) males tested individually. The solid line is the mean value at each time point, and the shaded region is the standard error of the mean.

### 2 Distance

The distribution of the Distance by Strain and Context is shown in Figure S2.1. MM and FM dyads had bimodal distributions, with a “close” peak around 3.3 mm (MM= 3.20, FM=3.39 mm, Fig. S2.1c,e) and a “distant” peak around 13 mm, which is close to the distance expected for flies that are randomly positioned within the arena. FF dyads only had the distant peak at 13.06 mm (Fig. 9a). The average distance in each context ± standard deviation was: FF=12.84 mm ± 0.96, FM=11.40 mm ± 2.11, MM=12.24 mm ± 1.51.

We assessed the effect of Context, Sex and Strain on Distance using ANOVA. We observed a significant effect of Context (F_2,1240_=114, p<2e-16, ω^2^=0.118, Figure S2.1a,c,e, Figure S2.2a) and Strain (F_4,1240_=56.7, p<2e-16, ω^2^=0.117, Figure S2.1b,d,f, S2.2b) and a significant interaction Context x Strain (F_8,1240_=26.6, ω^2^=0.107, Figure S2.1b,d,f, Figure S2.2c), while Sex and Sex x Strain were not significant, see Table S2.1 for the complete results. The significant Strain x Context interaction mainly reflects the fact that strains RAL-136 and RAL-338 had clear close peaks at about 3 mm in MM and FM dyads, while other strains lacked or showed reduced peaks there.

**Figure S2.1.**
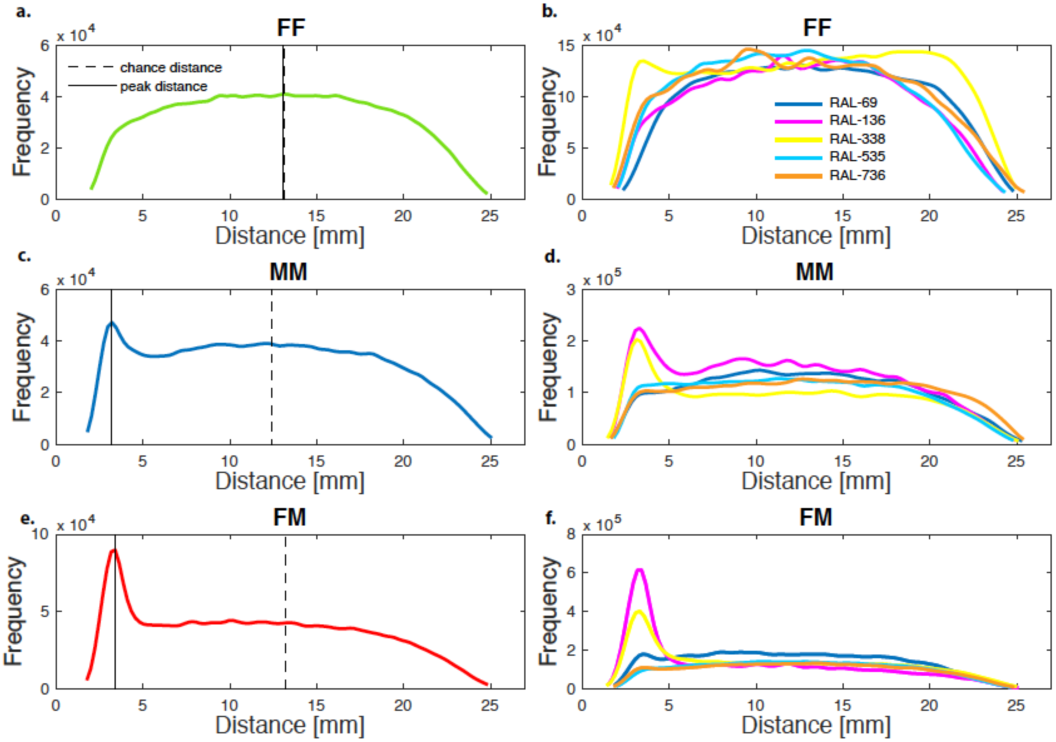
Distribution of Distance during the test in each Context and Strain (a) FF dyads, (b) FF dyads by strain, (c) MM dyads, (d) MM dyads by strain, (e) FM dyads, (f) FM dyads by strain. Dashed lines indicate the chance distance, black solid lines indicate the peak distance observed.

**Table S2.1.**
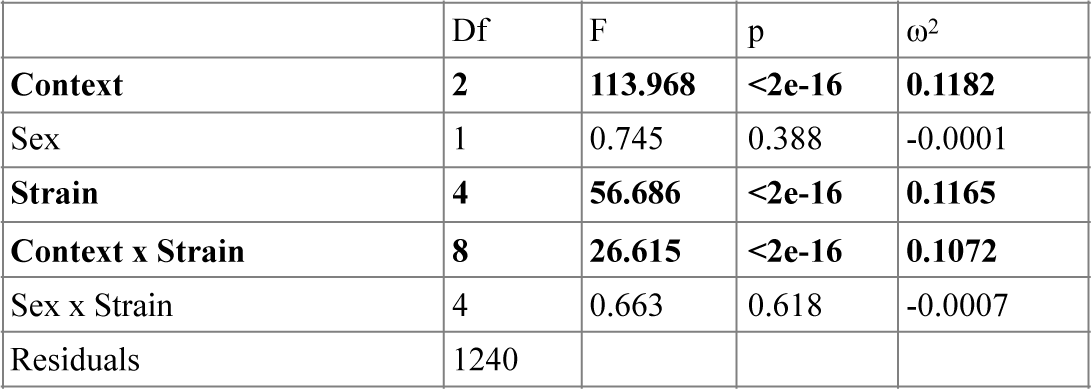
Results of the ANOVA and ω^2^ values on Distance with Context, Sex and Strain as independent variables. Bold cells indicate significant results.

| | Df | F | p | $\omega^2$ |
| --- | --- | --- | --- | --- |
| **Context** | **2** | **113.968** | **<2e-16** | **0.1182** |
| Sex | 1 | 0.745 | 0.388 | -0.0001 |
| **Strain** | **4** | **56.686** | **<2e-16** | **0.1165** |
| **Context x Strain** | **8** | **26.615** | **<2e-16** | **0.1072** |
| Sex x Strain | 4 | 0.663 | 0.618 | -0.0007 |
| Residuals | 1240 |  |  |  |

Post hoc ANOVAs showed significant effects of Context, but the effect size differed between comparisons: FF-FM (F_1,856_=159, p<2e-16, ω^2^=0.155), FF-MM (F_1,801_=53.1, p=7.7e-13, ω^2^=0.061), FM-MM (F_1,857_=39.1, p=6.33e-10, ω^2^=0.042). As expected from the interactions of courtship and mating, flies in the heterosexual dyads were closer than flies in the other contexts (Fig. S2.2a). Dyads with two females (FF) were significantly further than other dyads (Fig. S2.2a), about as far as expected from non-interacting flies. The fact that FM and MM dyads stayed closer than FF dyads, together with previous observations about intersex interactions (e.g., ^8^), is consistent with the males initiating courtship interactions in this species.

The distance between flies was clearly modulated by the social environment, genotype and their interaction. Flies in the FM and MM contexts stayed closer than expected by chance but only some genotypes (RAL-69, RAL-136 and RAL-338) had a distinct peak distance at around 3 mm. Previous studies have shown that flies communicate not only through chemosensory stimuli but also using mechanosensory interactions that induce flies to move in space ^4,9^. For this reason, genetic variation for distance has implications for the amount of information transferred between flies and their experience with the environment.

**Figure S2.2.**
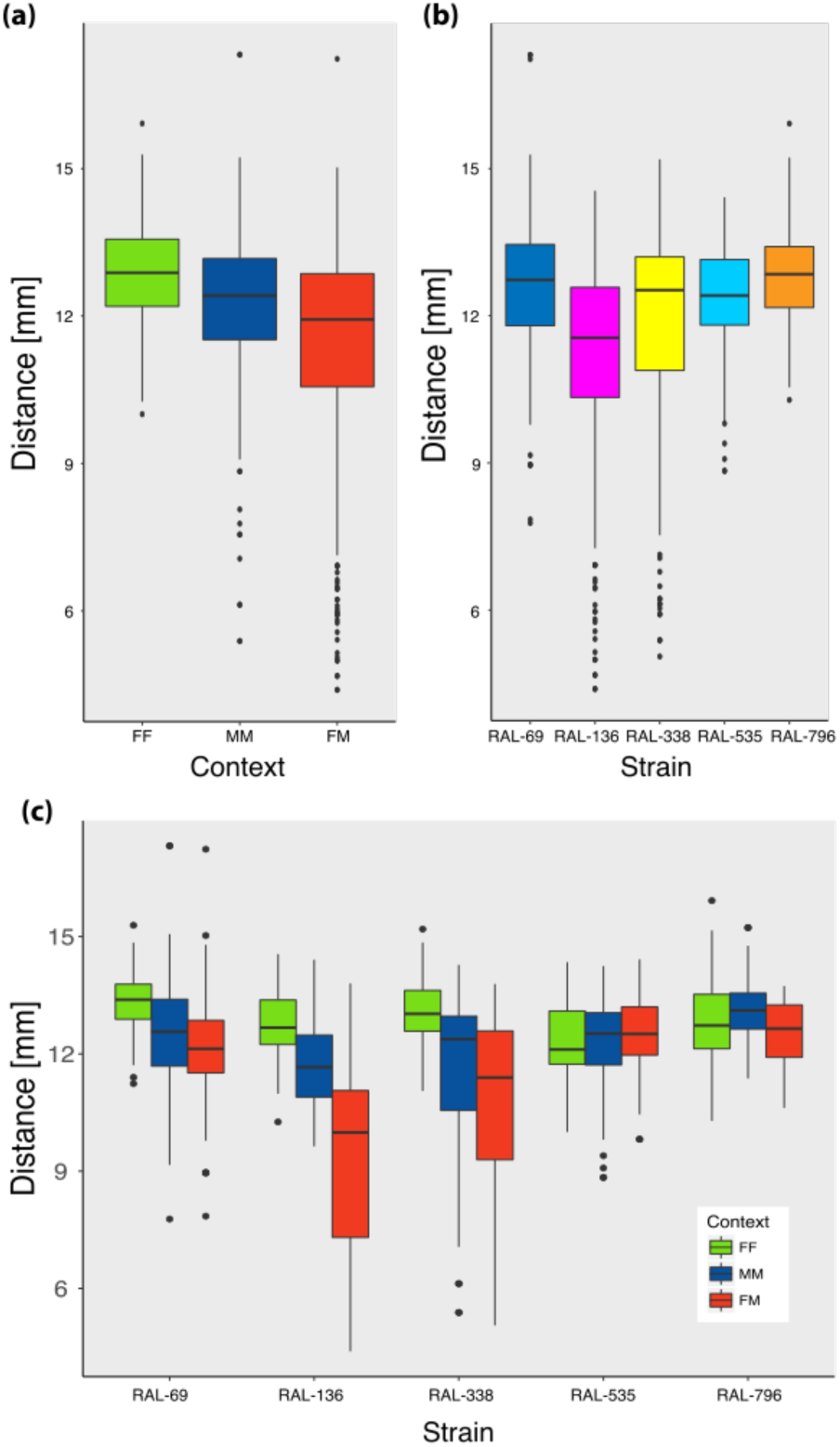
(a) Distance between flies by Context (FF, MM, FM), (b) by Strain (c) and by Context and Strain.

